# Improving behavior monitoring of free moving dairy cows using non invasive wireless EEG approach and digital signal processing techniques

**DOI:** 10.1101/2021.11.22.469585

**Authors:** Ana Carolina de Sousa Silva, Aldo Ivan Céspedes Arce, Hubert Luzdemio Arteaga Miñano, Gustavo Voltani von Atzingen, Valeria Cristina Rodrigues Sarnighausen, Ernane José Xavier Costa

## Abstract

**Background:** Electroencephalography (EEG) is the most common method to access brain information. Techniques to monitor and to extract brain signal characteristics in farm animals are not as developed as in humans and laboratory animals.

**New method:** The method comprised two steps. In the first step, the signals were acquired after the telemetric equipment was developed, the electrodes were positioned and fixed, the sample frequency was defined, the equipment was positioned, and artifacts and other acquisition problems were dealt with. Brain signals from six Holstein heifers that could move freely in free stalls were acquired. The control group consisted in the same number of bovines, contained in a climatic chamber (restrained group). In the second step, the signals were characterized by Power Spectral Density, Short-Time Fourier Transform, and Lempel-Ziv complexity.

**Results:** The results indicated that there was an ideal position to attach the electrodes to the front of the bovine’s head, so that longer artifact-free signal sections were acquired. The signals showed typical EEG frequency bands, like the bands found in humans. The Lempel-Ziv complexity values indicated that the bovine brain signals contained random and chaotic components. As expected, the signals acquired from the retained bovine group displayed sections with a larger number of artifacts.

**Comparison with existing methods:** We present the first method that helps to monitor and to extract brain signal features in unrestrained bovines.

**Conclusions:** The method could be applied to investigate changes in brain electrical activity during animal farming, to monitor brain activity related with animal behavior.

**Highlights:**

- A method that allows brain signals to be monitored in freely moving dairy cows is described
- The method uses noninvasive electrodes to minimize stress during EEG monitoring and allows bovines to behave normally during the process
- The method establishes the frequency sampling rate, electrodes positioning and fixation, equipment holding, artifact extraction, and signal characterization
- The brain signals are characterized by PSD, STFT, and Lempel-Ziv normalized complexity
- The method can be applied to relate EEG to animal behavior under normal handling conditions

## 1. Introduction

Animal behavior plays an important role in precision farming. Precision livestock farming is an approach that enables the farmer with more objective information about the animal to make better choices about the sustainability of their production system. Society in general is demanding closer attention to the needs of individual animal . Due to the increase in consciousness regarding animal welfare, the need for studies on animal behavior has begun to gain importance. In commercial livestock farming, the sustainability and profitability of production is ensured by raising the animals at the tolerable welfare level. The exposure of living organisms to conditions below the tolerable welfare level leads to the occurrence of physiological responses against stress conditions, resulting in increase in health problems and decrease in yield (Cam MA et al., 2018). In this scenario, understanding brain processes in bovines will allow some situations like heat stress (Silanikove, 2000; Sinha, 2007) and pain (McLean et al., 2017; Vuckovic et al., 2018), among others (Small et al., 2019), to be controlled.

Several DSP techniques have already been applied to humans and laboratory animals; e.g., FFT (Fast Fourier Transform) (Jalilifar et al., 2016), wavelets (Subasi et al., 2005), AGR (Adaptive Gaussian Representation) (Hashida et al., 2005), ANN (Artificial Neural Network) (Robert et al., 2002; Vuckovic et al., 2002), Lempel-Ziv complexity (Zhang and Roy, 1999), and entropy (Mazza et al., 2002), among others (Khosla et al., 2020; Motamedi-Fakhr et al., 2014). This kind of technology has also been employed to investigate some diseases in domestic pets (Bassett et al., 2014; Bergamasco et al., 2003; Brauer et al., 2012). Concerning farmed animals, studies involve pre-slaughter sensitization (Gibson et al., 2019; Small et al., 2019), sleep stages (Hänninen et al., 2008; Ternman et al., 2012) and pain (BERGAMASCO et al., 2011) most of them with animal kept in cages. This situation is changing and a recent study on sheep (Perentos et al., 2017), showed a method that did not use restraint and which employed implanted electrodes to analyze animal brain signals instead.

Acquiring brain signals in animals that can move freely presents various challenges, and the number of research studies dealing with bovine brain electrical activity is limited. Most of these studies have been conducted in laboratory conditions (Suzuki et al., 1990) and used subdermal electrodes (Jones et al., 1988; Suzuki et al., 1990) in animals that were subjected to some type of containment (Jones et al., 1988; Small et al., 2019). These conditions are unreal when we consider farmed animals, mainly large ones (Perentos et al., 2017).

Telemetric recording systems can monitor physiological parameters in freely moving animals without any restrictions in their exploratory behavior (Lapray et al., 2008). In 1966, West and Merrick (West and Merrick, 1966) presented a telemetric equipment for EEG acquisition in large animals and presented the test results obtained for cattle with their equipment. The authors showed the temporal characteristics of the signal, but they did not mention the characteristics in terms of frequency. After that, the need for long-term, continuous monitoring of EEG signals encouraged increasing use of telemetric systems, to provide continuous and reliable data for studies on epilepsy (Chang et al., 2011), sleep disorders (Weiergräber et al., 2005), and other pathologies (de Araujo Furtado et al., 2009) in freely moving animals. In almost five decades, different systems have been developed, from large systems (West and Merrick, 1966) to transmitters that can be implanted beneath the skin of small animals (e.g., rats) (Chang et al., 2011).

Another challenge regarding electroencephalography is the presence of artifacts in the signal. Due to the nature of the acquisition process, the EEG signal can mix with other biological potentials. The greatest influence is on the ocular signal (EOG), which is in the same amplitude range as the EEG (i.e., millivolts) (Urigüen and Garcia- Zapirain, 2015; Vidal et al., 2011), but it can still be contaminated by muscle (EMG) and cardiac (ECG) signals, both in the millivolt range (Reaz et al., 2006; Xie et al., 2021). Broadly speaking, artifacts can originate from internal (physiological activities of the subject) and external sources (environmental interference, equipment, electrode pop-up, and cable movement) and contaminate recordings in both temporal and spectral domains (Islam et al., 2016). When signals from animals are concerned, the presence of artifact is hardly avoided.

Given the number of research studies on bovine brain signal analysis and the kind of signal processing techniques they apply (Bager et al., 1990; Gibson et al., 2019; Jones et al., 1988; Merrick and Scharp, 1971; Small et al., 2019; Suzuki et al., 1990; Takeuchi et al., 1998; West and Merrick, 1966), a good strategy to characterize these signals is to assume that they are nonstationary, as in humans, and to employ time– frequency techniques to estimate the spectra.

EEG is typically described in terms of rhythms and transients. The rhythmic activity of EEG is divided into frequency bands (Islam et al., 2016; Ramadan and Vasilakos, 2017). This characteristic makes traditional analysis rely mainly on the detection of spectral power changes (Derya Übeyli, 2009). The Power Spectral Density (PSD) is calculated by Fourier Transform, the estimated autocorrelation sequence that is found by nonparametric methods. One of these methods is Welchs’s method (Al- Fahoum and Al-Fraihat, 2014). Because the EEG signal is nonstationary, the most suitable way to extract feature from raw data is to use time–frequency domain methods, such as STFT (Short-Time Fourier Transform) (Kıymık et al., 2005; Marchant, 2003; Ramos-Aguilar et al., 2020). STFT is a time-dependent Fourier Transform that may be calculated by sliding a window over the signal to compute the FFT (Marchant, 2003). It is represented by a two-dimensional time–frequency plot, called spectrogram. The major drawback of STFT is a compromise between time and frequency resolution. If a short window is applied, information derived from each FFT will be well localized in time, but frequency resolution will be poor.

In addition, the electrical activity of the brain (EEG) exhibits significant complex behavior with strong nonlinear and dynamic properties (Jeong, 2004; Zhang et al., 2001). Non-linearity in the brain is introduced even at the cellular level because the dynamic behavior of individual neurons is governed by threshold and saturation phenomena (Abásolo et al., 2006). This implies that nonlinear methods can be applied to investigate EEG dynamics (Zhang et al., 2001).The Lempel-Ziv complexity (LZC) (Lempel and Ziv, 1976) uses symbolic techniques to map a time series into a sequence that retains its dynamics. LZC has been widely used in biomedical applications to estimate the complexity of discrete-time signals (Gómez et al., 2006). Normalized complexity can be used to quantify nonlinear and nondeterministic data (Costa et al., 2017) .

This study aimed to develop a method to monitor bovine brain electrical activity by noninvasive techniques in animals in an experimental free stall. Another objective was to apply techniques like PSD (Power Spectral Density), STFT, and LZC to characterize the obtained signals, a necessary step to expand this field of study.

## 2. Material and Methods

This section is divided into three subsections: (1) developing a portable and low-cost wireless electroencephalograph; (2) developing a method to monitor bovine brain electrical activity; and (3) extracting the features of these signals.

### 2.1. Wireless electroencephalograph

The EEG acquisition system consisted of two modules: the first module, which was embedded in the animal, concerned the recording, conditioning, and wireless transmission of the EEG signals. The second module, which was called base module, received data from the embedded module and connected the system to a computer (Silva et al., 2005).

#### 2.1.1. Animal module

This module could be attached to the animal’s neck, and it performed three tasks: (1) signal amplification and conditioning, (2) analog-to-digital conversion, and (3) wireless transmission of the digital data. A microcontroller sampled the EEG analog signal, converted the analog signal to a digital signal, and transmitted the recorded data (Silva et al., 2005). Each of these parts will be detailed below.

The electronic amplification and conditioning circuit consisted of (i) an amplifier that pre-amplified the EEG signal and (ii) three analog active filters: a 0.05-Hz high-pass filter, a 1.5-kHz low-pass filter for antialiasing, and a band-pass filter to remove 60-Hz electromagnetic interference. The total gain of the circuit was 1 x 10^4^.

The A/D converter stage of the circuit used a 10-bit analog-to-digital converter of a Microchip PIC16F877A microprocessor and allowed frequency up to 120 Hz to be sampled.

Through its USART interface, the microprocessor also controlled telemetric transmission. The circuit used the BIM2-433-160 transceiver (Radiometrix Ltd, 2004), which transmitted and received the data at 433 MHz with low energy consumption in a range of up to 200 m.

#### 2.1.2. Base module

The base module received and stored data arriving from the embedded modules. This module consisted of one 433-MHz channel that used a BIM2 transceiver (Radiometrix Ltd, 2004). Dedicated software in the computer stored the data in a database and showed the EEG in real-time on the display monitor.

#### 2.1.3. Equipment validation

To validate the test equipment (TE), the TE ability to amplify signals of low amplitude and frequency was evaluated and compared to the control equipment (CE), which consisted of a 32-channel digital electroencephalograph from the company EMSA Medical Equipment S/A, model BRAINNET BNT-EEG. The following signals were evaluated: (1) 3.90-Hz and 30.20-Hz sine waves generated by a sine generator and (2) brain signals from a 26-year-old volunteer free of neurological problems. Stretches of 5 and 10 s were collected at a sampling frequency of 100 Hz for sine waves and brain signals, respectively.

In the frequency domain, the PSD was estimated by the Welch method for each signal. In the time domain, the comparison was made by considering the Signal-to-Error Ratio (SER) (Equation 02). The method considered the developed equipment as the output of a nonlinear predictor of the EEG signal acquired by the control equipment (Kavitha and Narayana Dutt, 1999).

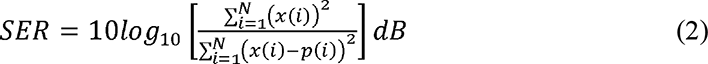

### 2.2. Signal monitoring

To develop a method to monitor bovine brain electrical activity, it was necessary to define sample frequency (Fs), where to place the electrodes (positioning and fixation) and equipment, and to deal with artifacts and other acquisition issues. These conditions were established by conducting measurements with two animals. The second stage involved 12 animals, which were divided into two groups of six, and brain signals were acquired in two different conditions: from animals moving freely in a free stall and from animals kept in a climatic chamber.

Considering that the frequency interval of human brain signals ranges from 0.4 Hz to almost 40 Hz (Islam et al., 2016; Ramadan and Vasilakos, 2017), Fs was initially set to 100 Hz, to ensure that frequencies were mapped between 0 and 50 Hz without aliasing. This allowed the presence of frequencies higher than the frequencies detected in humans to be investigated. The signals were digitized in the embedded module.

The experiment was carried out in the city of Pirassununga, state of São Paulo, Brazil (latitude 21°59’46” south and longitude 47°25’33” west, altitude of 627 meters). The signals were acquired from Holstein heifers.

This study was conducted by using noninvasive electrodes consisting of 1.0-cm gold discs. Electrode fixation was attempted with either strong adhesive tapes or cyanoacrylate, also known as super glue (Super bonder - Loctite®). First, the area at the forehead of the cows where the electrodes would be fixed was shaved. Then, the electrodes were filled with conductive gel for EEG. Next, the animal module was placed in a small bag around the animal’s neck. Fixation was tested for a whole month.

The equipment comprised a single-channel electroencephalograph (bipolar measurement), so the sensor consisted of two electrodes, to acquire the biopotential signal acquisition, and a grounding electrode. The grounding electrode was positioned at the back of the animal’s neck, whereas the other two electrodes were tested at three different positions (POS1, POS2, and POS3) of the animal’s forehead. These positions were selected on the basis of the bovine head anatomy. Anatomical head sections were analyzed to investigate positions for electrode fixation. The images of heads that were used for this analysis were obtained at a local slaughterhouse, so they are not the images of heads of the animals that were used in the experiments.

To minimize artefacts from eyes and muscle movements, the test was performed on animals sedated with 1 mL/kg Rompun from Bayer®. A total of 6 mL was enough to achieve deep sedation for 40 min. The positioning test data were acquired from two cows during a whole day.

After the best electrode position was determined, the signals were acquired in two different situations: 1) from animals contained in a climatic chamber (cage containment) and 2) from animals in pasture (animals with freedom of movement). In the first situation, the animals were contained in 1.0 x 2.0 m individual cages arranged as a 2x3 matrix inside a climatic chamber. In the second situation, the animals were contained in free stalls measuring 4.0 x 3.0 m and arranged side by side as a 1x6 matrix. The cows had free access to food and water during the experiment. For each situation, the signals were acquired from six Holstein heifers.

During the experiment, the temperature and relative air humidity were collected and used to determine the enthalpy of the place, as proposed by Albright (1990). The enthalpy is a variable that is considered an index of thermal comfort—it indicates the environmental conditions related to the thermal stress suffered by the animals (Moura, D. J.; Naas, I. A.; Silva, I. J. O.; Sevegnani, 1997; Rodrigues et al., 2011). This stage of data acquisition lasted three full days.

### 2.3. Signal characterization

For both groups of animals, all the stages of brain signal characterization were performed with the Signal Processing Toolbox^TM^ from MATLAB^®^ software releases R2015a and 2019a, developed by The Mathworks, Inc.

In pre-processing stages, different methods can be employed to handle artifacts.

One of them rejects the epoch or segment of EEG data that is labelled as artifactual (Motamedi-Fakhr et al., 2014). Sauter et al. (Sauter et al., 1990) proposed a sequential method where spectral analysis is performed after corrupted segments are extracted. For this purpose, artifacts are detected by visual inspection, and then rejected. Welch’s method was applied to artifact-free signal segments to estimate the signal PSD and to verify whether the spectral content is due to brain signal.

PSD makes the frequency spectrum smoother than the raw FFT output, and the Welch algorithm (Monson H. Hayes, 1996) is a nonparametric method to estimate PSD. Let us consider that the FFT is calculated over the entire duration of the signal. In the Welch algorithm, instead of processing the FFT over the entire time domain, the signal is separated in windows with the same size. Window size affects the clarity of the result by cutting frequencies with periods larger than the window. Windowing is taking a sample of a larger dataset and tapering the signal at the edges of each interval. This makes the signal smoother without sharp transitions that can disturb the frequency spectrum representation (Same et al., 2020). Welch’s method was applied to artifact- free signal segments to estimate the signal PSD.

Considering that STFT is a sequence of Fourier Transforms of a windowed signal, it can be represented as in Equation (3) (Kehtarnavaz, 2008).

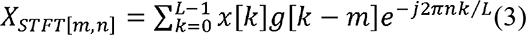

where x[k] denotes the signal, and g[k] denotes an L-point window function.

STFT was also employed to artifact-free signal segments, so that data could be mapped in the time–frequency domain.

For LZC calculation, each signal segment is converted into a binary sequence s(n), as follows (Bachmann et al., 2018; Gómez et al., 2006):

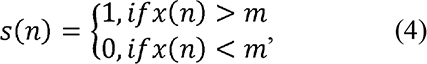

where x(n) is the signal segment, and n is the segment sample index from 1 to N (segment size). The segment length N was chosen for at least 400 samples – a value for which normalized LZC is stabilized (Gómez et al., 2006); m is the signal mean value. Thereafter, the resulting binary sequence s(n) is scanned from left to right, and the number of different patterns is counted. The complexity value c(n) is increased every time a new pattern is encountered (Bachmann et al., 2018; Gómez et al., 2006; Lempel and Ziv, 1976). Lempel and Ziv (Lempel and Ziv, 1976) showed that

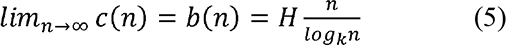

where N is the segment length, k is the number of different symbols in the sequence (in the binary case, k = 2, and H represents entropy (Equation (6)).

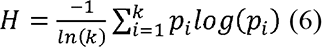

where p indicates the probability that a state i is obtained by counting the occurrences of each symbol, divided by the total number of symbols in the sequence.

To avoid the variations due to the segment length, normalized LZC values are calculated as follows:

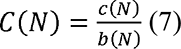

The more random the sequence, the greater the number of different patterns present in it; that is, its normalized complexity C(N) approaches one. Meanwhile, C(N) values that approach zero are associated with deterministic sequences. Normalized LZV - C(N) was applied to artifact-free signal segments to investigate oscillatory, chaotic, and random components in bovine brain signals. To illustrate the relationship between C(N) and signal characteristics, low- and high-complexity known sequences were calculated.

## 3. Results and Discussion

### 3.1. Equipment validation

Figure 1 presents the graph of a 3.90-Hz sine wave sampled at 100 Hz by the test equipment (TE) and control equipment (CE). In this case, SER was 39.35 dB.

**Figure 1.**
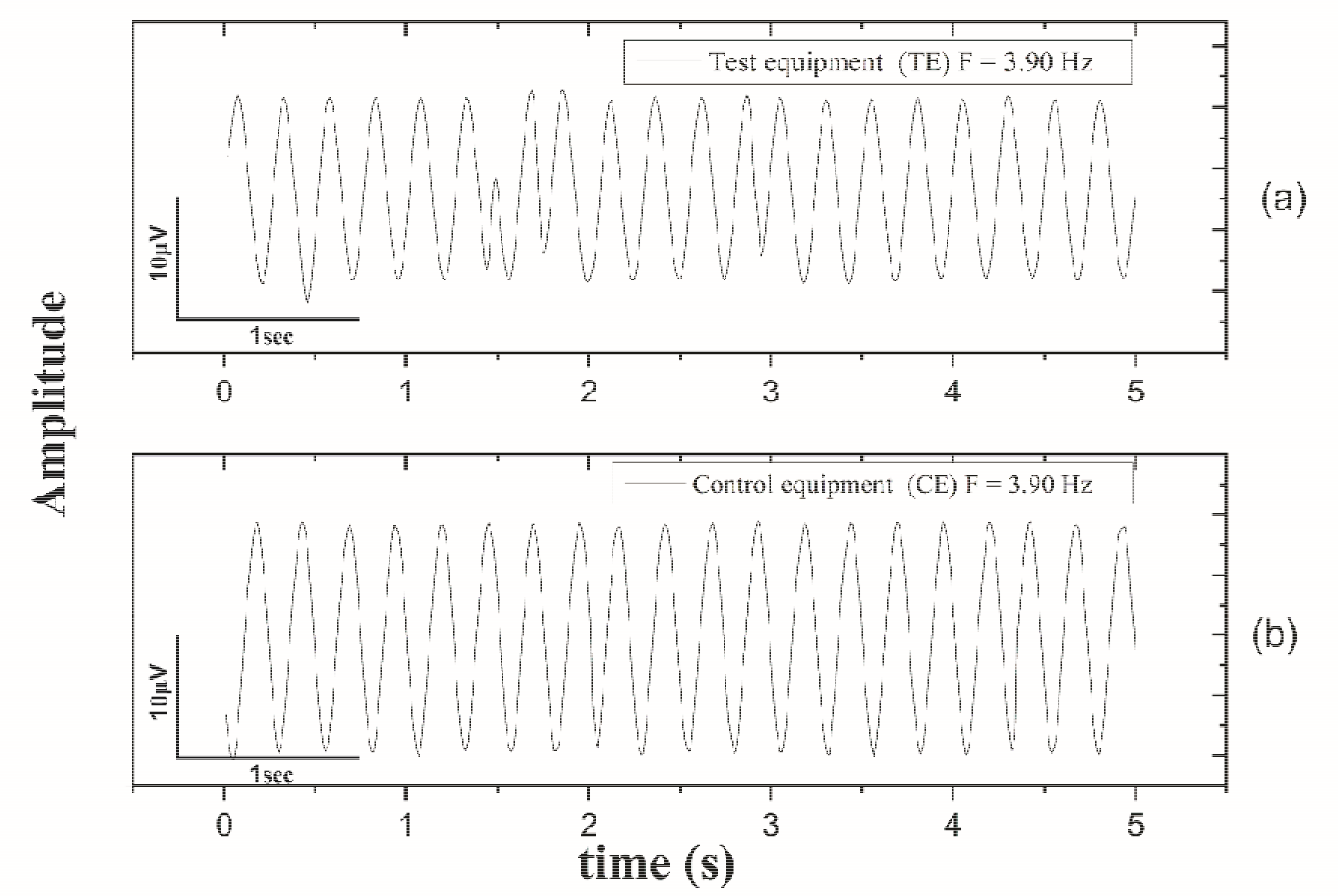
3.90-Hz sine wave sampled at 100 Hz by (a) test equipment (TE) and (b) control equipment (CE).

Figure 2 shows the graph of a 30.2-Hz sine wave sampled at 100 Hz by the test equipment (TE) and control equipment (CE). In this case, SER was 38.36 dB.

**Figure 2.**
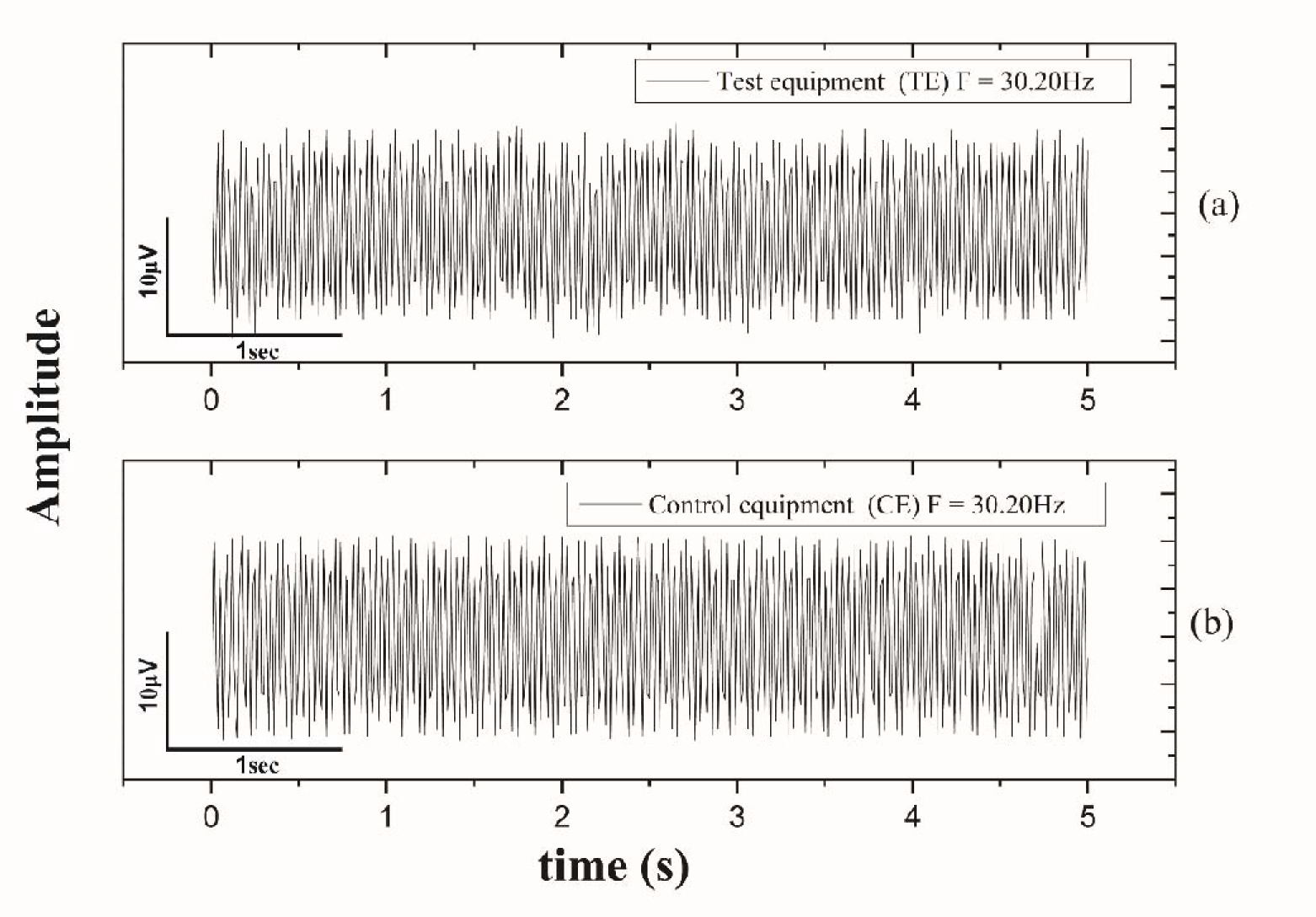
30.2-Hz sine wave sampled at 100 Hz by (a) test equipment (TE) and (b) control equipment (CE).

Figure 3 compares the PSD of both sine waves obtained with the TE and CE.

**Figure 3.**
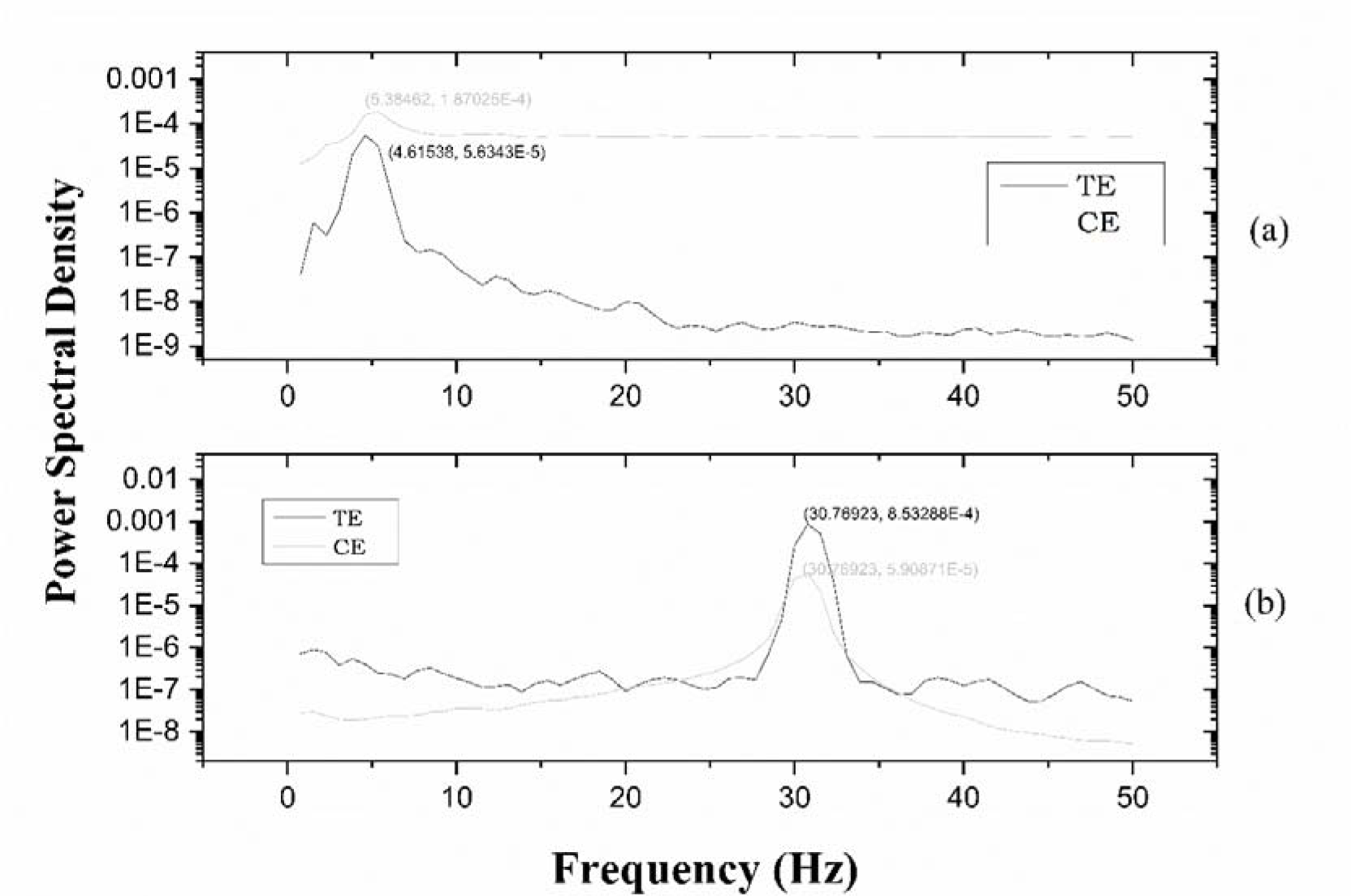
Power spectral density (PSD). (a) 3.90-Hz sine wave and (b) 30.2-Hz sine wave. Test equipment (TE) in black and control equipment (CE) in gray.

Figure 4 compares the brain signals acquired from a human volunteer by both the TE and CE. Figure 5 illustrates the PSD calculation for the signals in Figure 4. In this case, SER was 2.41 dB.

**Figure 4.**
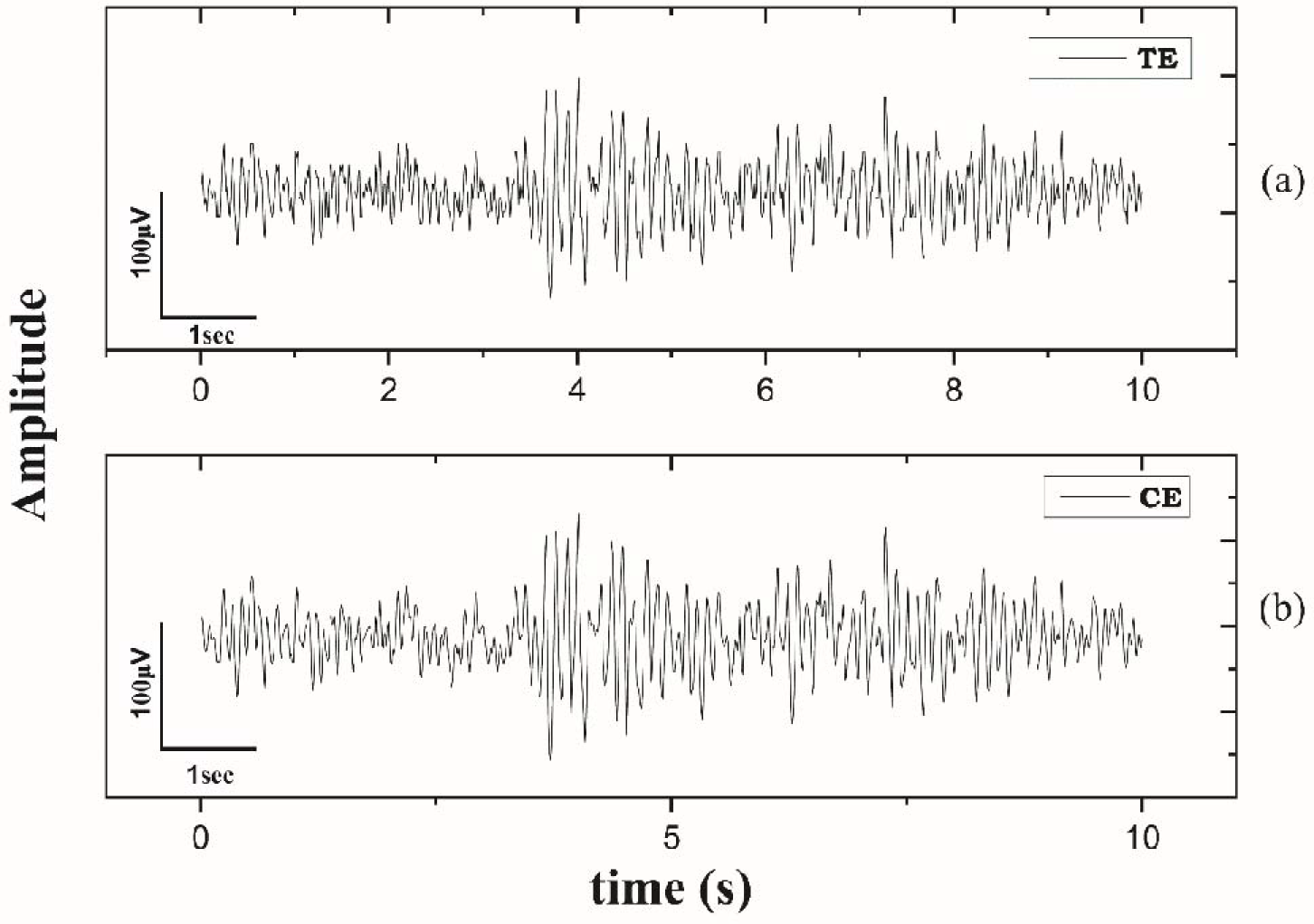
EEG of a 26-year-old female volunteer recorded with (a) test equipment (TE) and (b) control equipment (CE).

**Figure 5.**
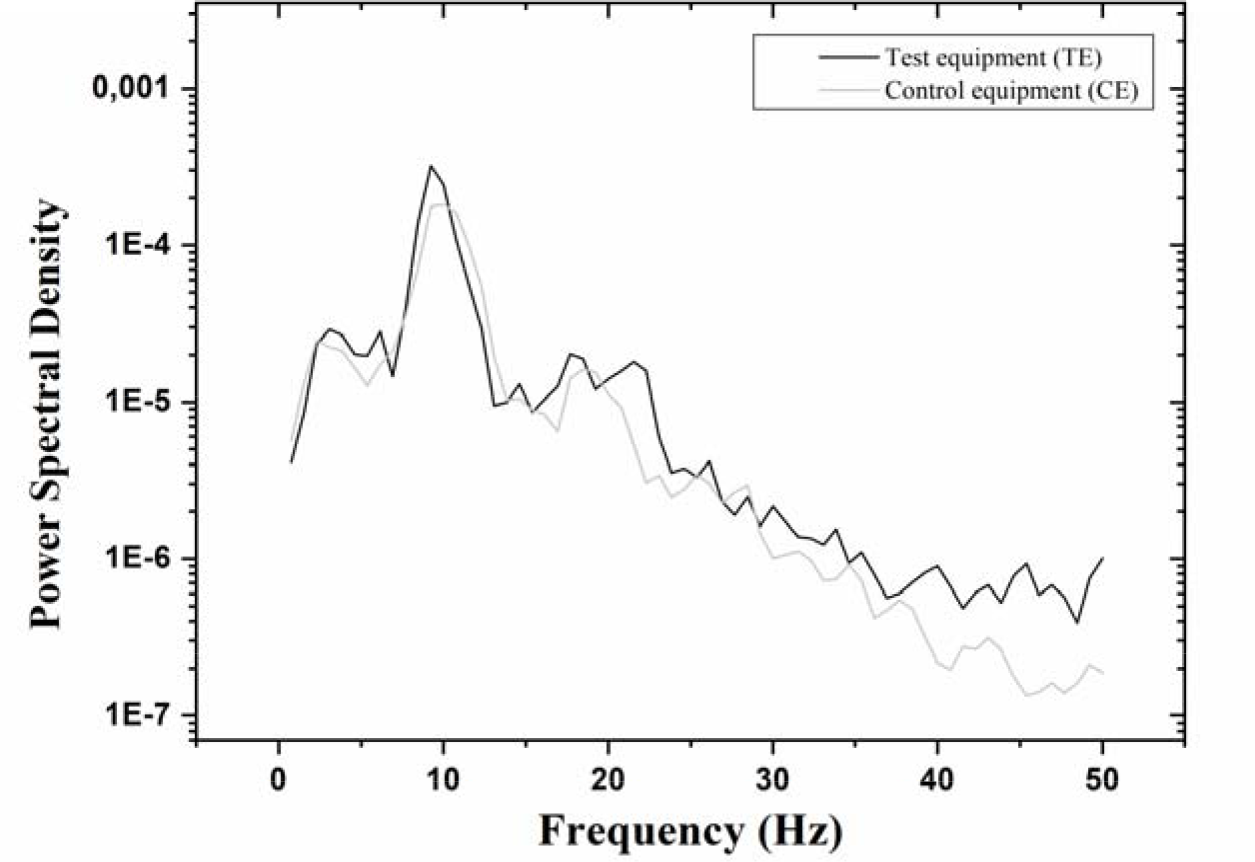
Power spectral density (PSD) for the human volunteer. Test equipment (TE) in black and control equipment (CE) in gray.

In the time domain, the signals were only slightly significantly different as compared to the signals obtained with the method proposed by Kavitha and Narayana Dutt (1999). The main difference between the signals was due to equipment resolution, which was 10 and 12 bits for the TE and CE, respectively. The PSD indicated the same frequencies for both the generated sines and brain signals.

The first telemetric equipment developed to record EEG (West and Merrick, 1966) weighed approximately 0.450 kg, transmitted data up to 91.44 m, and was completely analog. The equipment cost about US $1500, including the receiver. The literature has no other report on the development and use of other EEG equipment for use in cattle.

The equipment developed herein weighs 180 g and transmits data up to 200 m. In addition, it has a microcontrolled processing system that allows digital data transmission and costs approximately US$ 100.

### 3.2 Signal monitoring

Figure 6-a shows a head section of an adult bovine. This figure illustrates the distance from the brain to the frontal region of the bovine head. This distance can be longer or shorter, depending on the paranasal sinuses. Finding the best position for signal acquisition involves locating the region where this distance is as short as possible.

**Figure 6.**
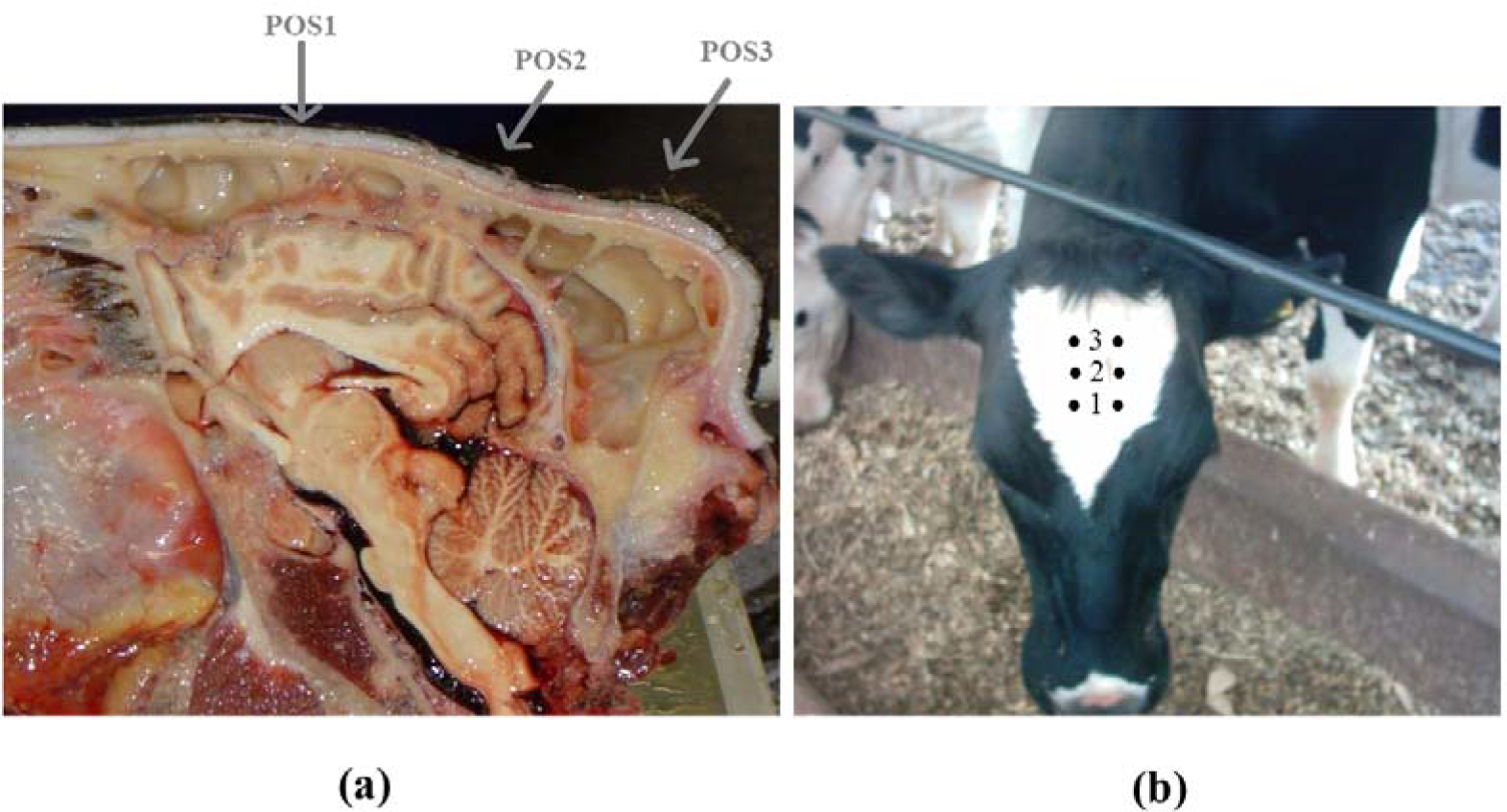
(a) Head section from adult animal. POS2 corresponds to the region where the brain is closest to the surface. (b) Layout of monitoring electrodes at three positions. Numbers 1, 2, and 3 indicate POS1, POS2, and POS3, respectively.

Figure 6-b shows the layout of the monitoring electrodes at the three evaluated positions.

Figure 7 depicts a disc electrode and electrode fixation with adhesive tape.

**Figure 7.**
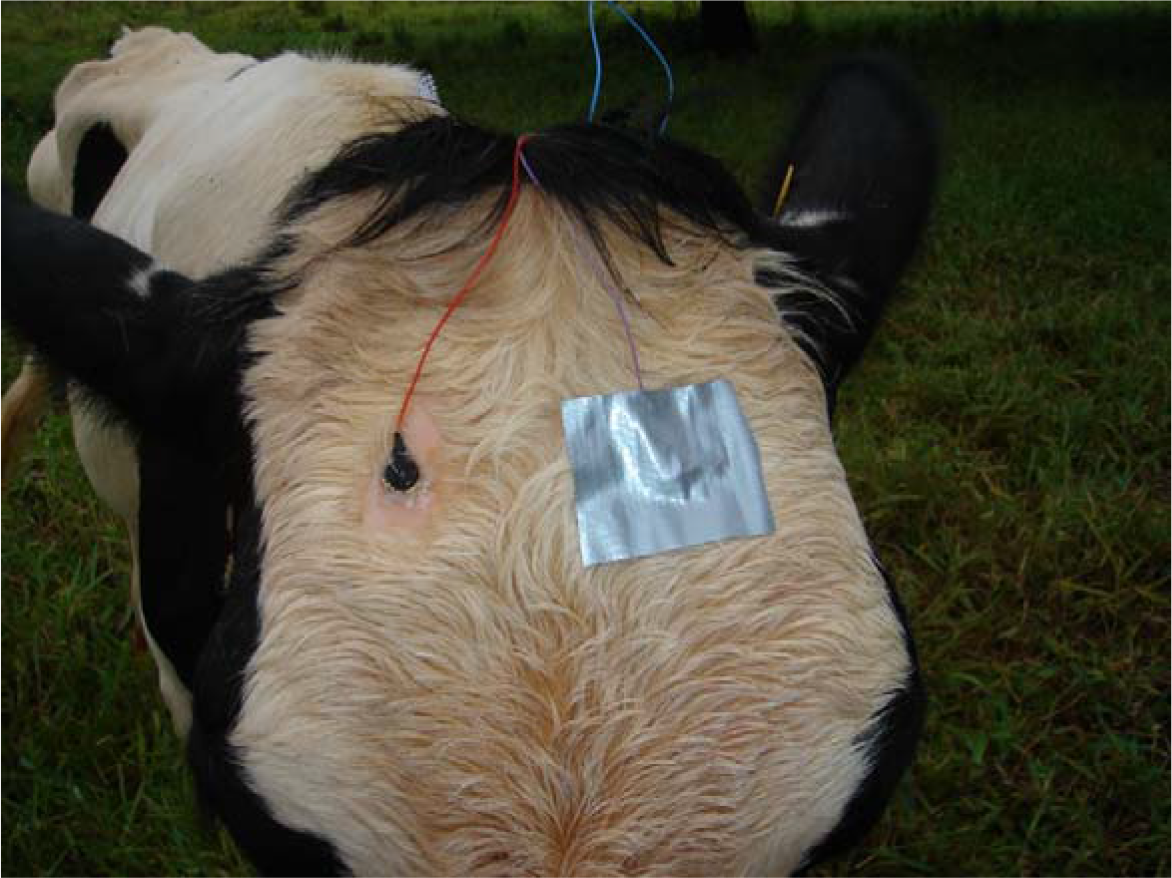
Disc electrodes at POS2.

The animal module consisted of a box measuring 2.5 x 7.0 x 11.0 cm^3^ and weighing 180 g. To make it less uncomfortable for animals, the box was placed in a fabric bag, which was attached to the neck with a soft ribbon. Figure 8-a presents the animal module placed at the animal’s neck, whilst Figure 8-b shows the halter that was used to protect the electrodes and the electrodes fixed with super glue.

**Figure 8.**
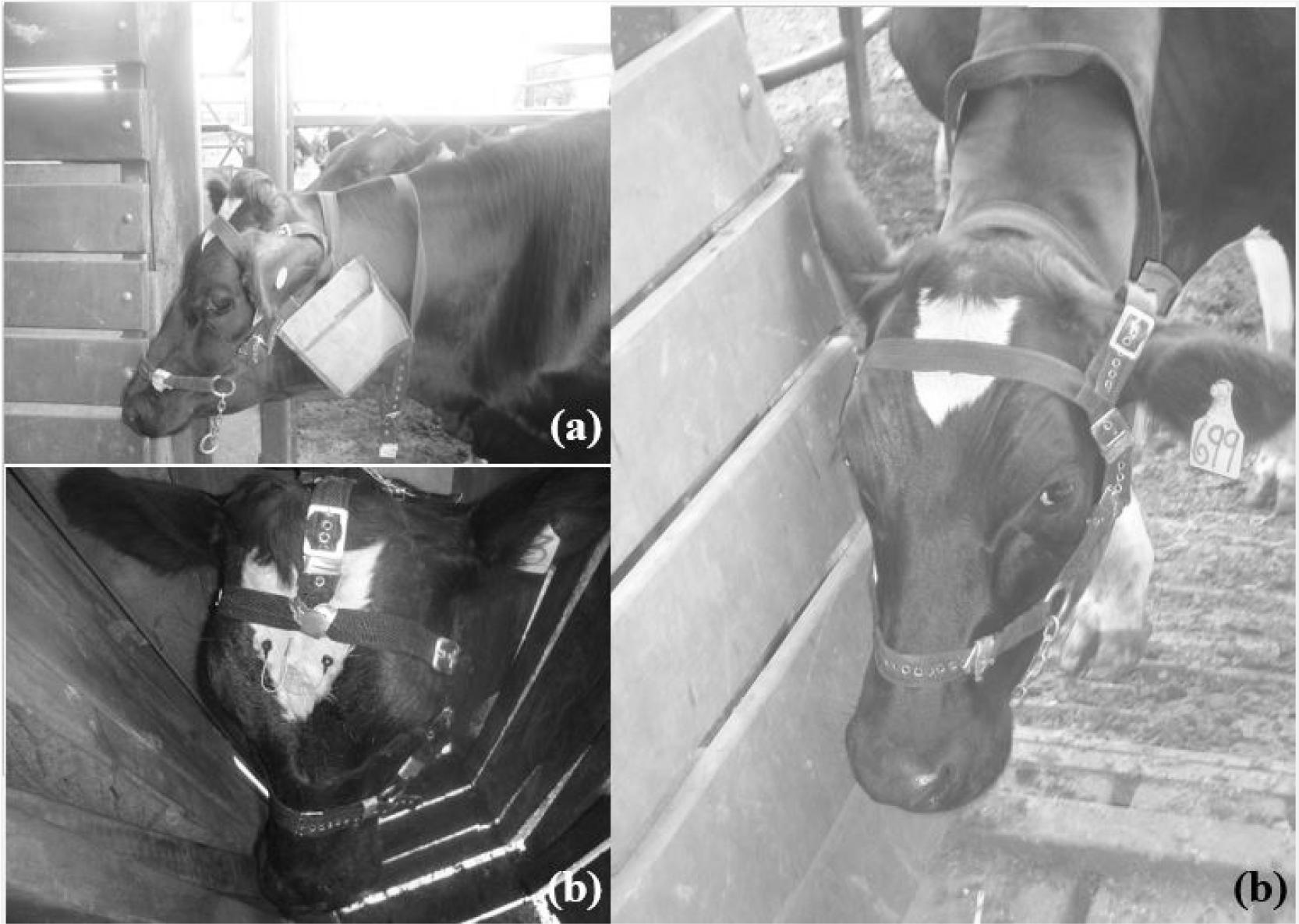
(a) Animal module inside a fabric bag placed at animal’s neck. (b) glue fixation of electrodes and protection with halter.

The adhesive tape failed to keep the electrodes rigidly attached to the animal, so they often came loose. The glue (Figure 8-b) kept the electrodes in the right position for a whole day of acquisition. At the end of the day, the electrodes and the animal’s forehead had to be cleaned for a new acquisition. Compared to the use of implanted electrodes, which can go on for a year (Perentos et al., 2017), this is a short period. It is important to highlight the possibility of noninvasive monitoring here, though.

Figure 9 contains signals acquired at the three selected positions for one of the two animals in this stage of the experiment, while Figure 10 corresponds to the artifact- free signal sections in Figure 9.

**Figure 9.**
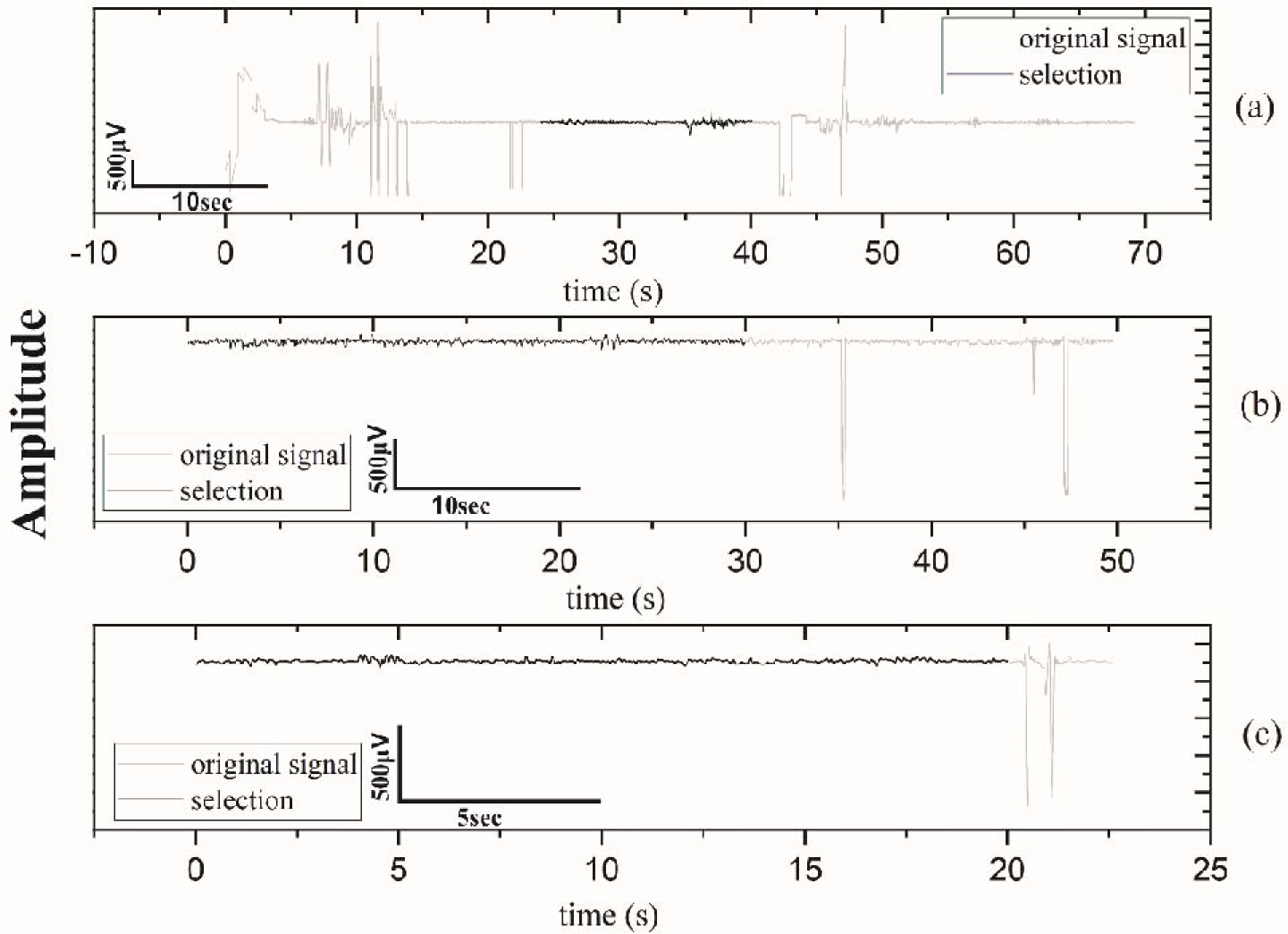
Signal at (a) POS1. (b) POS2. (c) POS3. Original signal in grey and selected signal epochs in black.

**Figure 10.**
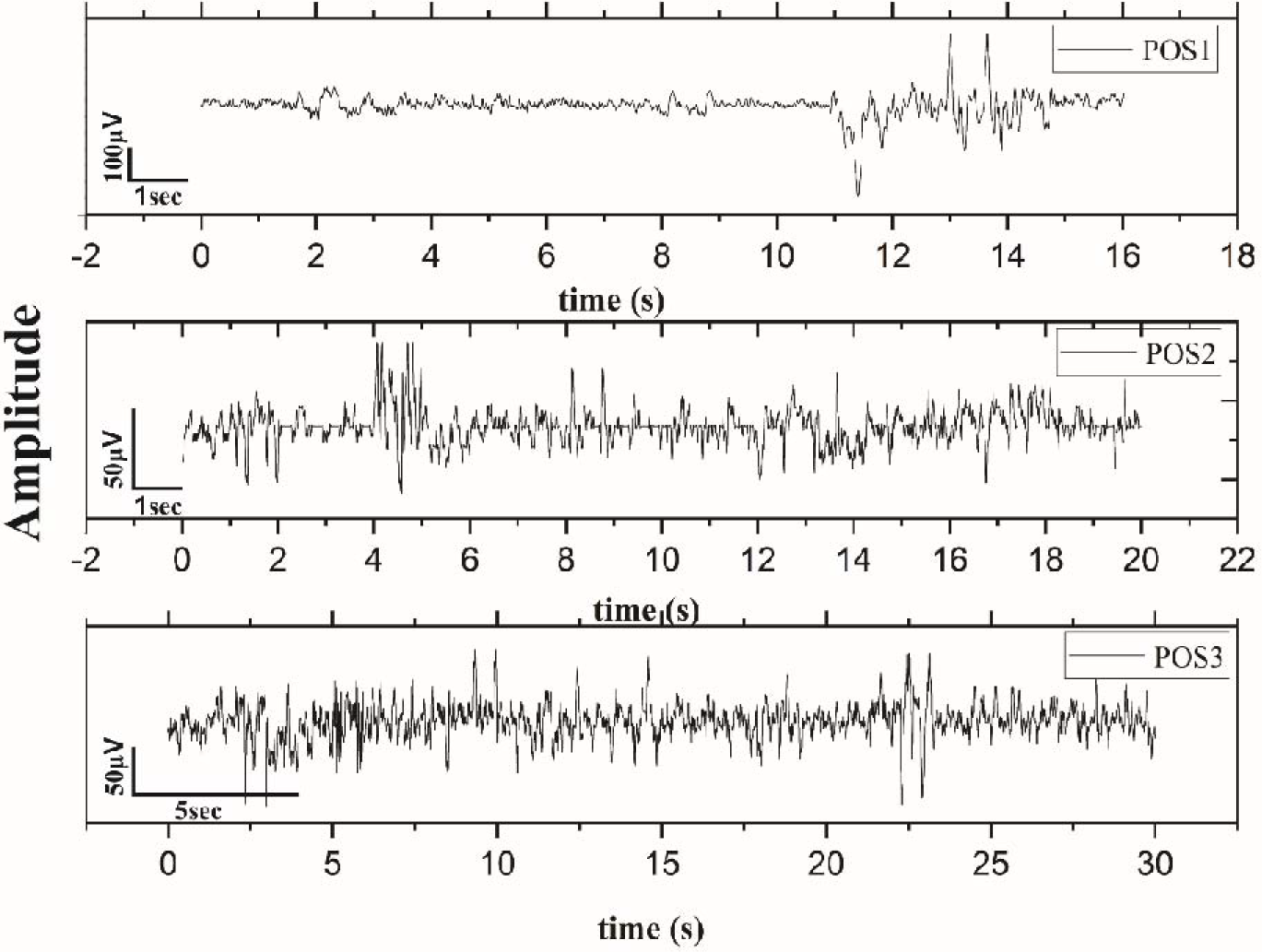
Artefact-free signal sections in (a) POS1, (b) POS2, and (c) POS3.

Figure 11 contains the PSD for the signal sections in Figure 10.

**Figure 11.**
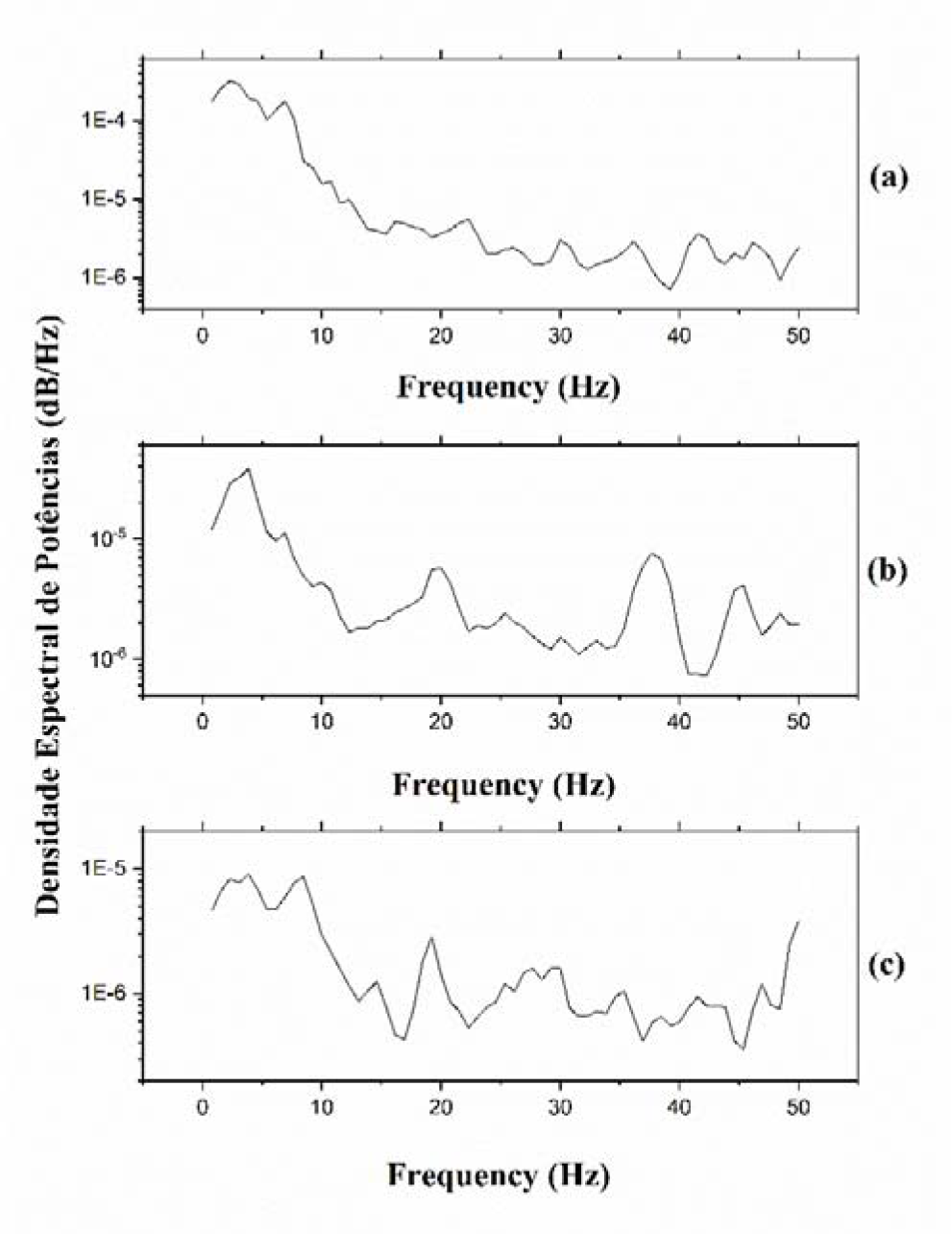
PSD for artifact-free signal sections at (a) POS1, (b) POS2, and (c) POS3.

In Figure 9-a, the signal at POS1 presented many artifacts, most of which were due to ear movement. POS2 provided longer artifact-free sections than POS1 and POS3 as a result of two main facts. First, POS3 was largely influenced by eye movement, whereas POS2 is the region where paranasal sinuses were smaller and hence the surface was closer to the cerebral cortex. This result was expected given the analyzed anatomical head section (e.g., Figure 6). The three positions had similar PSD profile, and the observed frequencies (2–10Hz; 13–27Hz) agreed with those obtained by Suzuki et al (1990)(Suzuki et al., 1990) and Ternman et al (2012). Therefore, the metric to choose the best electrode position was the length of the obtained artifact free-section, and POS2 was selected as the best one to acquire brain signals in cattle.

Most studies in this area have been conducted in laboratory conditions (Suzuki et al., 1990) and used subdermal electrodes (Jones et al., 1988; Suzuki et al., 1990) in animals that were subjected to some type of containment (Jones et al., 1988; Small et al., 2019). The method presented herein allows brain signal to be recorded with surface electrodes in freely moving animals.

Figure 12 displays enthalpy values for the animals in climatic chamber or pasture during the first of the three days of experiment.

**Figure 12.**
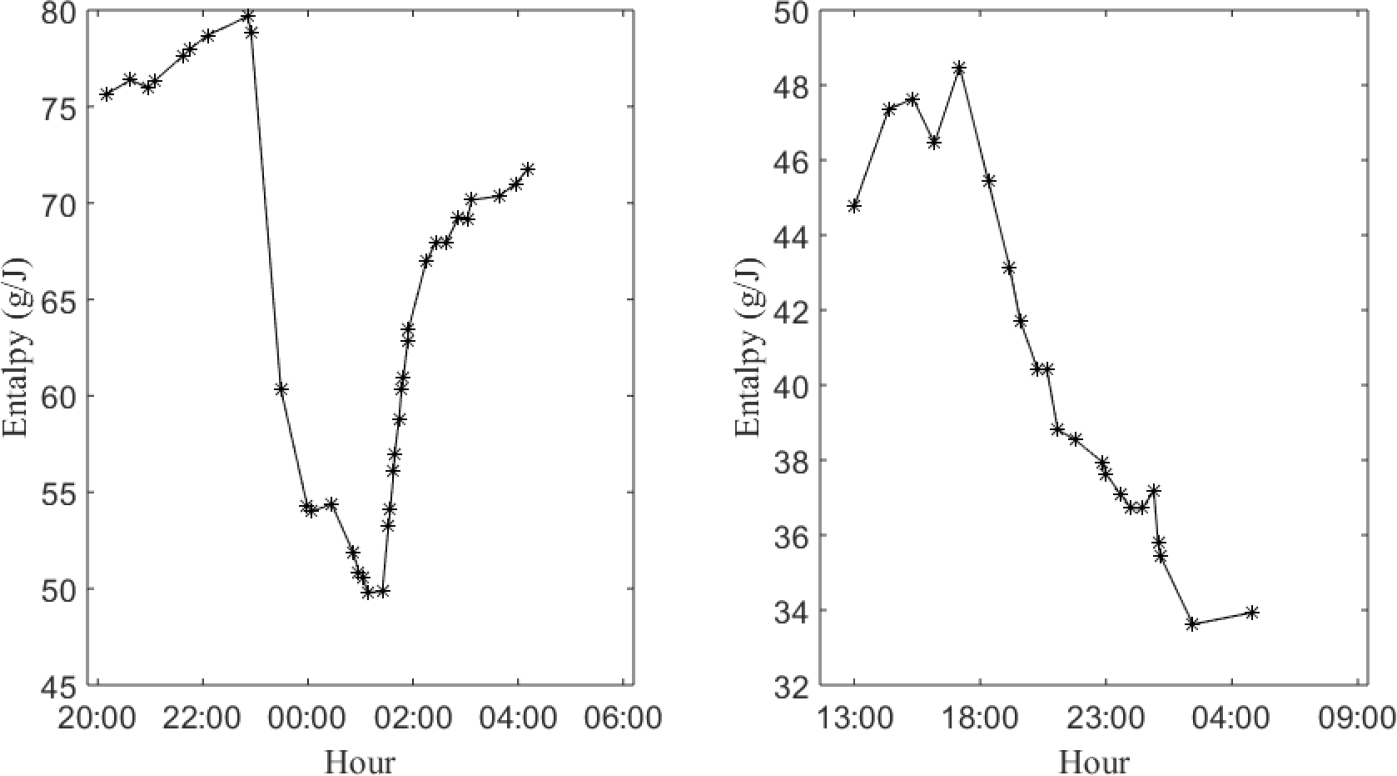
Enthalpy for animals in (a) climatic chamber and (b) pasture.

In the climatic chamber, between 20:00 h and approximately 22:30 h and after 03:00 h, the animals experienced a heat stress situation, but most of the time, they were in a comfortable situation (30–70 J/kg of dry air).

The stress condition was due to animal agitation in periods of enthalpy above 70 J/g, which provided a greater amount of water vapor resulting from animal panting. This increased the internal relative humidity, which progressively became an obstacle to heat loss through breathing and the skin surface. Under these conditions, the temperature and relative air humidity were above 32 °C and 60%, respectively. After a period of reduction in animal agitation and renewal of the chamber air, the condition of thermal comfort was established, with average values of 25 °C and 60 %. In the pasture, there were no conditions of thermal stress. In fact, during the analysis period, enthalpy decrease, as the incidence of solar radiation decreased, which modified the temperature and relative air humidity (Rodrigues et al., 2011).

### 3.3 Signal characterization

Figure 13 exemplifies a pre-processing stage when artifact was removed from the signal for one of the animals in the climatic chamber (13-a) and one unrestrained animal (13- b).

**Figure 13.**
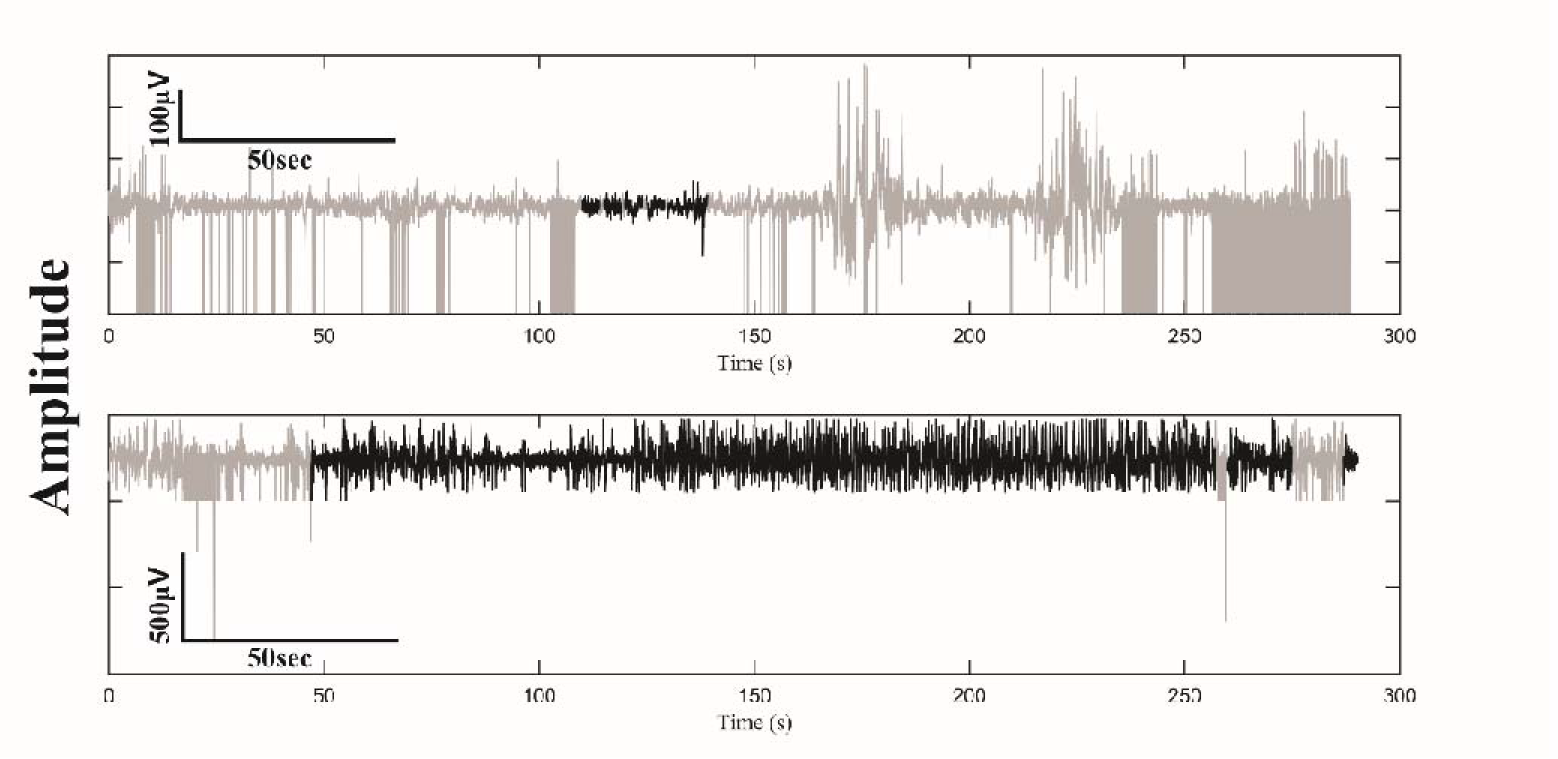
Artifact removal. (a) Climatic chamber. (b) Pasture. Original signal in grey and signal selected epochs in black.

The proportion of artifact-free data in relation to the original data seen in Figure 13 was observed for most of the database. In all the acquisitions, at least one artifact-free section could be selected.

Figure 14-a contains the PSD for the artifact-free signal section in Figure 13-a, while Figure 14-b shows the PSD for the second artifact-free signal segment in Figure 13-b.

**Figure 14.**
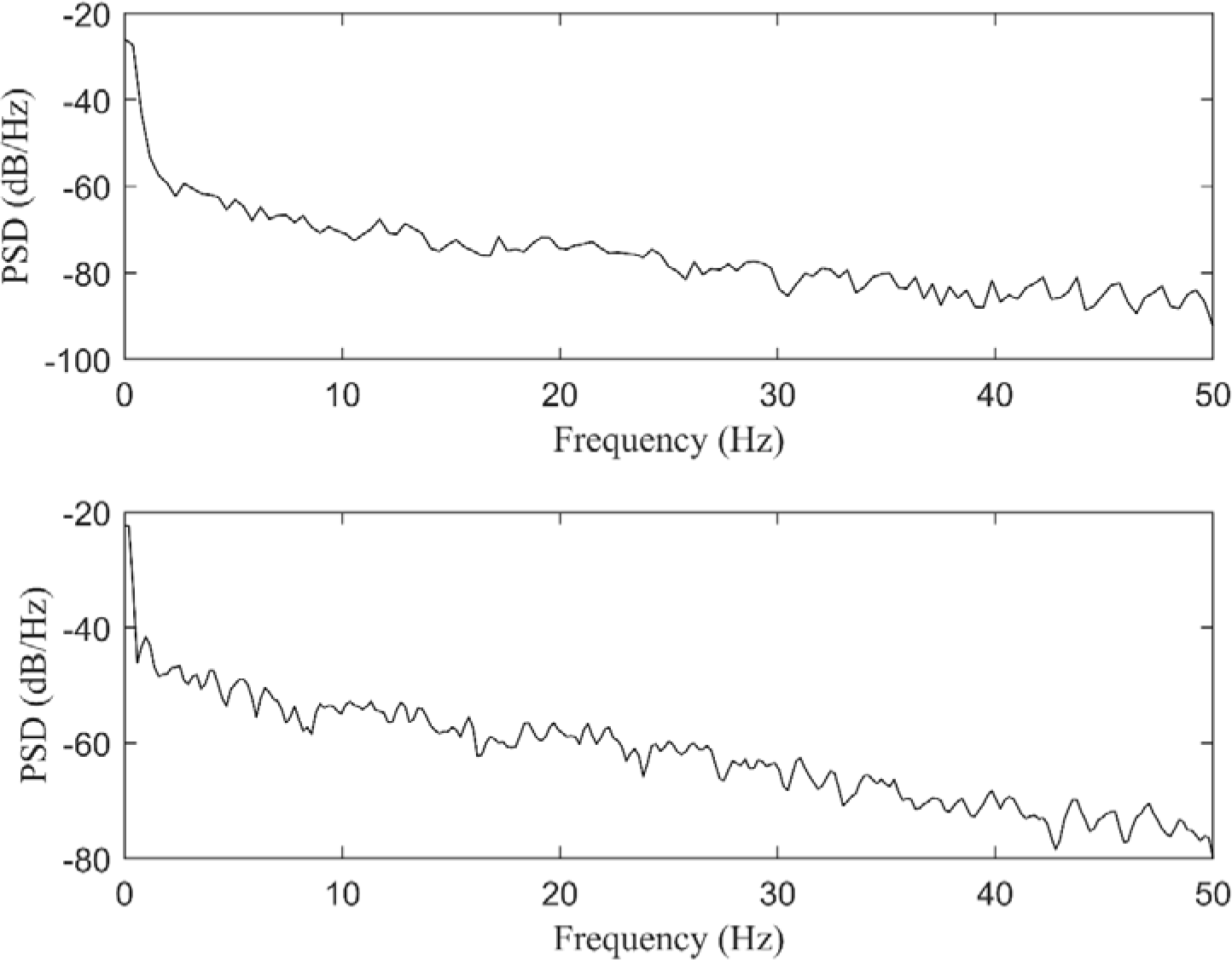
Power Spectral Density (PSD) for artifact-free EEG signal. (a) Climatic chamber. (b) In pasture.

We were not able to establish a direct relationship between enthalpy and any specific behavior observed during the experiments (eating, drinking water, chewing, standing, lying down, or walking). The stress experienced by animals in the cage overlapped with any other behavior. This reflected even on the duration of artifact-free epochs in the signal (less than 4 s for all the animals).

Figure 15 illustrates STFT for the second artifact-free signal segment in Figure 13-b.

**Figure 15.**
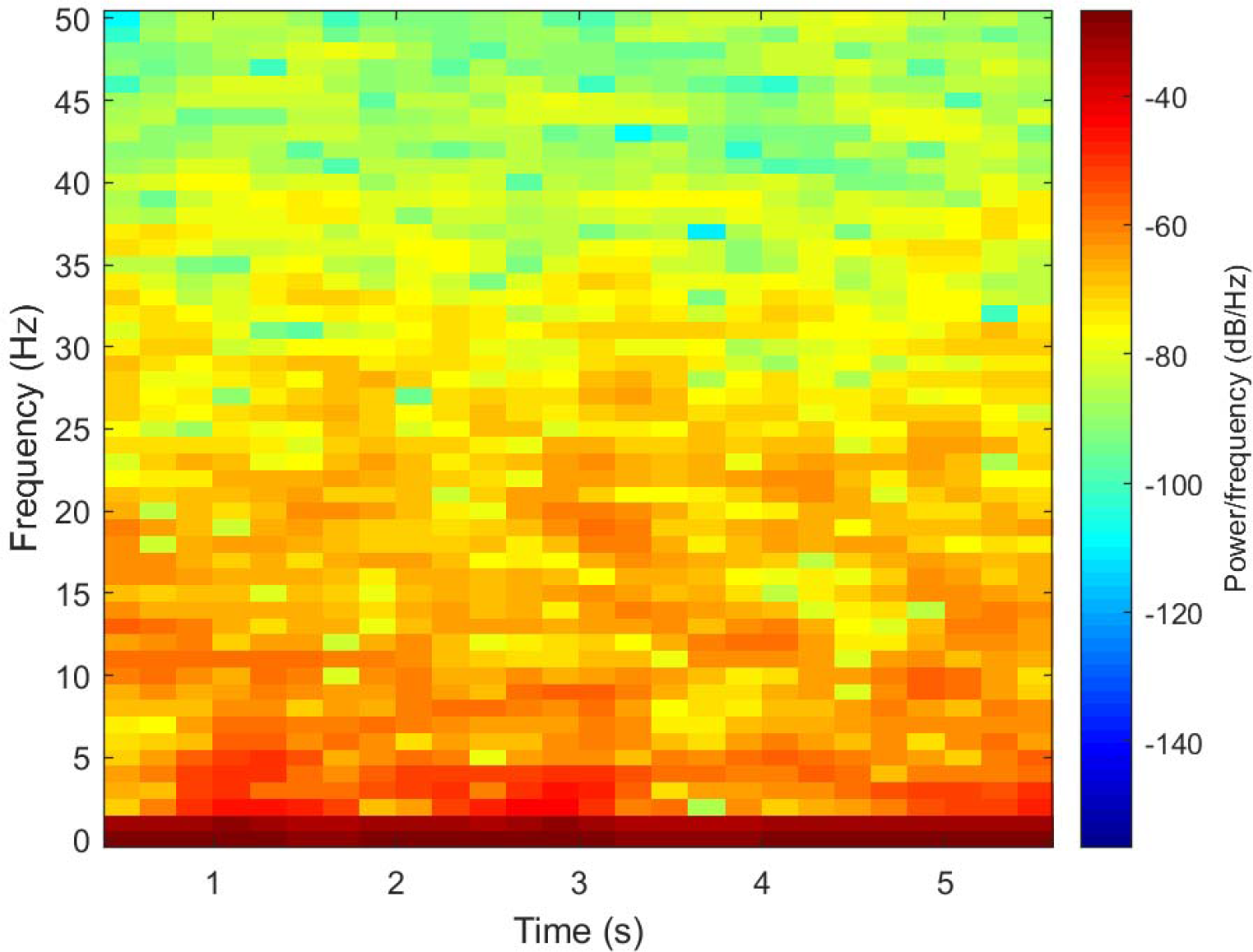
STFT for the second artifact-free EEG signal segment in Figure 13-b.

We were not able to associate the observed frequencies in the group with any specific animal behavior, such as eating, drinking water, chewing, standing, lying down or walking, mainly because of the amount of data that was acquired. However, the spectrogram clearly showed the predominant frequencies in cattle signals (Figure 15). As expected, the delta wave presented high amplitude between 1 and 3 Hz and was very expressive between 1 and 2 Hz. There were also well-delimited regions between 4 and 7 Hz (Theta), 7 and 12 Hz, (Alpha), 12 and 30 Hz (Beta), and above 30 Hz (Gamma).

Observed frequencies still agreed with the observations of Suzuki et al. (1990) and Takeuchi et al. (1998).

This methodology allowed us to acquire brain signals from animals that can move freely while minimizing the stress generated by confinement. It also allowed the acquisition to occur noninvasively. However, the acquisition conducted in this way produced data with a greater number of artifacts, but it had the advantage of reducing the discomfort caused by using subdermal electrodes. Another important point to consider is the challenge imposed by the size of the animals. Considering the larger amount of data, this methodology could be applied to investigate changes in brain electrical activity during animal farming, to monitor brain pathologies and other situations.

Table 1 lists normalized complexity for seven knowledge sequences.

**Table 1.** Lempel & Ziv normalized complexity C(N) for different time series.

| Sequence | Cn |
| --- | --- |
| $S1 = [0\ 1\ 0\ 1\ 0\ 1\ 0\ 1\ 0\ \dots 1\ 0\ 1\ 0\ 1\ 0\ 1\ 0\ 1\ 0\ 1]$ | 0.0202 |
| $S2 = \sin(2\pi 30t)$ | 0.0559 |
| $S3 = \sin(2\pi 0.5t) + \sin(2\pi 10t) + \sin(2\pi 18t) + \sin(2\pi 30t) + \sin(2\pi 30t)$ | 0.0129 |
| $S4 = \sin(2\pi 0.5t) + \sin(2\pi 10t) + \sin(2\pi 18t) + \sin(2\pi 30t) + \sin(2\pi 30t) +$<br>(array of random numbers) | 1.0000 |
| $S5 = \text{white noise}$ | 0.9957 |
| $S6 = \text{array of random numbers}$ | 1.0870 |
| $S7 = \text{logistic map in chaos threshold value (Costa et al., 2017)}$ | 1.0000 |

Table 2 contains Cn for artefact-free signal sections in Figure 13.

**Table 2.** Normalized complexity Cn for artefact-free signal sections

| Sequence | Cn |
| --- | --- |
| Climatic chamber 1<sup>st</sup> section (size: 4 s) | 0.9894 |
| Climatic chamber 2<sup>nd</sup> section (size: 4 s) | 0.6596 |
| Climatic chamber 3<sup>rd</sup> section (size: 4 s) | 0.7483 |
| Climatic chamber 4<sup>th</sup> section (size: 4 s) | 0.7475 |
| Climatic chamber 5<sup>th</sup> section (size: 9 s) | 0.5857 |
| Pasture 1<sup>st</sup> section (size: 240 s) | 0.6078 |
| Pasture 2<sup>nd</sup> section (size: 15 s) | 0.6163 |
| Pasture 3<sup>rd</sup> section (size: 3 s) | 0.8656 |

We obtained values similar to the values in Table 2 for the other five animals we evaluated. High normalized LZC C(N) values can be associated with the presence of noise, random numbers, or the predominance of one of these events (Table 1). On the basis of the values we observed for bovine signals (Table 2), we can consider that the artifacts were not completely eliminated, and that cattle brain signals have some random characteristics. Costa et al. (2017) used the logistic map in the chaotic behavior threshold to obtain the entropic q-index for different values of initial conditions, to find that C(N) always presented values around one. Thus, complexity values next to one might indicate that bovine brain signals also have chaotic components.

Future improvements to this method could include increasing the number of animals and intensifying signal acquisition during a specific behavior (eating, drinking water, chewing, standing, lying down, or walking), so that said behavior could be associated with signal features. Another point for the future is to handle artifacts automatically, given that this is a time-consuming stage of signal processing.

## 5. Conclusions

EEG signals can be acquired from unrestrained cattle by using surface (noninvasive) electrodes and telemetric equipment. There is an ideal position for electrode attachment to the front of the head, so that longer artifact-free signal sections can be acquired. The signal presents typical EEG frequency bands, such as the bands found in humans and other animals. The signals also display random and chaotic characteristics. As expected, the restrained animals present signals with more artifacts. The method developed herein can be applied to understand several situations involving bovine (or other bigger farm animals) brain signals ranging from normality to some brain pathologies.

## Funding

This work was supported by the Sao Paulo State Research Foundation [Grant Number 03045911]

## Author contributions

**Ana Carolina de Sousa Silva**: Conceptualization, Methodology, Investigation, Data curation, Formal analysis, Writing- Original draft preparation, Writing- Reviewing and Editing. **Aldo Ivan Céspedes Arce**: Software, Data curation, Project administration, Writing- Original draft preparation. **Hubert Luzdemio Arteaga Miñano**: Formal analysis. **Gustavo Voltani von Atzingen**: Formal analysis, Writing- Reviewing and Editing: **Valeria Cristina Rodrigues Sarnighausen**: Investigation, Writing- Reviewing and Editing. **Ernane José Xavier Costa**: Conceptualization, Methodology, Investigation, Writing- Reviewing and Editing, Project administration, Supervision, Funding.

## Abbreviations

AGR: Adaptive Gaussian Representation
EEG: Electroencephalogram
LZC: Lempel and Ziv complexity
C(N): Lempel& Ziv normalized complexity
DSP: Digital Signal Processing
FFT: Fast Fourier Transform
Fs: Sample frequency
POS1: Electrodes evaluation position 1
POS2: Electrodes evaluation position 2
POS3: Electrodes evaluation position 3
PSD: Power spectral density
STFT: Short-Time Fourier Transform
EOG: Electroocoulogram
ECG: Electrocardiogram

## Notes

### Competing Interest Statement

The authors have declared no competing interest.

